# *In Vitro* Analysis of Bacterial Microcompartments and Shell Protein Superstructures by Confocal Microscopy

**DOI:** 10.1101/2022.08.24.505129

**Authors:** Daniel S. Trettel, Wade C. Winkler

## Abstract

The shell proteins that comprise bacterial microcompartments (BMCs) can self-assemble into an array of superstructures such as nanotubes, flat sheets, and icosahedrons. Physical characterization of BMCs and these superstructures typically relies on electron microscopy, which decouples samples from their solution context. We hypothesize that an investigation of fluorescently tagged BMCs and shell protein superstructures *in vitro* with high resolution confocal microscopy will lead to new insights into the solution behavior of these entities. We find that confocal imaging is able to capture nanotubes and sheets previously reported by TEM. Using a combination of fluorescent tags, we present qualitative evidence that these structures intermix with one another in a hetero- and homotypic fashion. Complete BMCs are also able to accomplish intermixing as evidenced by colocalization data. Finally, a simple colocalization experiment suggests that fluorescently modified encapsulation peptides (EPs) may prefer certain shell protein binding partners. Together, these data demonstrate that high resolution confocal microscopy is a powerful tool for investigating microcompartment-related structures *in vitro*, particularly for colocalization analyses. These results also support the notion that BMCs may intermix protein components, presumably from the outer shell.

**Importance:** Microcompartments are large, organelle-like structures that help bacteria catabolize targeted metabolites, while also protecting the cytosol against highly reactive metabolic intermediates. Their protein shell self-assembles into a polyhedral structure of approximately 100-200 nm in diameter. Inside the shell are thousands of copies of cargo enzymes, which are responsible for a specific metabolic pathway. While different approaches have revealed high-resolution structures of individual microcompartment proteins, it is less clear how these factors self-assemble to form the full native structure. In this study, we show that laser scanning confocal microscopy can be used to study microcompartment proteins. We find that this approach allows researchers to investigate the interactions and potential exchange of shell protein subunits in solution. From this, we conclude that confocal microscopy offers advantages for studying the in vitro structures of other microcompartments, as well as carboxysomes and other bacterial organelles.

## Introduction

Bacterial microcompartments (BMCs) are a widely distributed class of prokaryotic organelle. ^1, 2, 3^ These sub-cellular structures are typically 40-300 nm in diameter and composed entirely of protein. The protein elements are segregated into a distinct icosahedral protein shell and its inner enzymatic cargo. ^1, 4^ The inner cargo is responsible for a diverse range of chemistries,^2, 3, 5^ from carbon fixation^6^ to glycyl-radical enzyme mechanisms.^7, 8^ The overall modular arrangement of BMCs has led researchers to propose repurposing BMCs into an accessible platform for synthetic biology efforts.^4, 9, 10, 11^ However, synthetic adaptations of BMCs have been generally slowed by inadequate knowledge of the “rules” that govern BMC assembly.

The BMC protein shells self-assemble into a honeycomb-like lattice that selectively gates the influx and efflux of key metabolites. Specifically, the facets of the shells are made from a combination of flat, cyclic homo-hexamers (BMC-H) and homo-trimers (BMC-T), and the vertices are capped by similarly arranged pentamers (BMC-P).^12, 13^ Recently, high-resolution cryo-electron microscopy experiments have captured exquisite detail of synthetic BMC shells, providing important clues into the way the shell “tiles” piece together.^14, 15, 16, 17^ While these synthetic shells are small (typically <40 nm in diameter), larger BMC shell platforms have also been reported.^18, 19^

Individual shell proteins are not capable of polymerizing into the polyhedral BMC structure; instead, combinations of different classes of shell proteins are thought to be required for this to occur. This implies that there are intrinsic physicochemical properties of the different classes of shell proteins that guide their successful polymerization into polyhedral structures. Interestingly, in the absence of the correct partners, some of the individual shell proteins have been reported to form alternate super-structures. For example, a few BMC-H proteins have been noted to self-assemble into tube-like structures (coined ‘nanotubes’) when purified *in vitro*.^20, 21, 22, 23, 24^ Nanotubes have also been reported for certain BMC-T proteins as well,^21^ although BMC-P types have not been associated with these structures. These purified nanotubes were observed by electron microscopy in bundles or as individual tubes. ^21, 24^ Bundles of BMC-H nanotubes may even be capable of forming within bacterial cells upon overexpression of the BMC-H proteins.^19, 20, 23, 24^ Further, these BMC-H nanotube bundles appear to interfere with septation, thereby resulting in elongated and linked cells.^23^

These nanotube structures have not been systematically investigated or analyzed by solution-based experimental approaches. It is not known whether they are still capable of interacting with other shell proteins or with cargo proteins. Therefore, a deeper investigation of these remarkable structures will give new and important insight into the protein-protein interactions that occur within BMC. It would also potentially lead to new synthetic biology tools.^25^ For example, it is tempting to imagine adapted versions of these shell proteins being used as synthetic cytoskeletal elements. And, by extension, it is not inconceivable that certain BMC shell proteins may have evolved to function as natural cytoskeletal features. Simply put, it is unclear how these higher-order structures behave in solution, whether they can interact with other BMC proteins, and if they are biologically relevant. For all these reasons, the super-structures formed by purified BMC shell proteins warrant considerable further study.

Herein, we apply multiple imaging techniques to investigate the structures formed by BMC shell proteins *in vitro*. Specifically, we implemented solution-based confocal techniques to visualize BMC-related structures. Such methods open new avenues for studying solution properties of these structures. We chose to analyze well-characterized shell proteins used by the *Salmonella enterica* propanediol utilization (Pdu) BMC. To analyze the proteins and their associated structures, we developed an approach wherein these structures, including native BMCs, could be visualized by laser scanning confocal microscopy. As a solution technique, confocal microscopy further allowed colocalization events to be observed when different shell proteins are equilibrated, suggesting intermixing of components. When preparations of BMCs that had been tagged with different fluorophores were mixed, we found that these fluorophore-tagged proteins also exhibited increasing colocalization over time. Consistent with this, we find that native BMC can associate with the nanotubes. Finally, we showcase another utility of this mode of analysis to directly show that encapsulation peptides (EP), which are known to target cargo protein to the inner lumen of BMC, associate with certain shell protein structures. Together, our data gives proof of principle evidence that confocal microscopy-based approaches can be used to study BMC-related structures in solution, including their propensity to exchange proteins subunits.

## Results

### The hexamer PduA forms nanotubes that can be imaged with confocal microscopy

BMCs and nanotubes formed from BMC shell proteins have been previously investigated by transmission electron microscopy (TEM).^21, 23, 24^ While TEM remains a powerful tool for analyzing BMC-related structures, we reasoned that BMC proteins might also be imaged using laser scanning confocal microscopy if the proteins were fluorescently labeled. While suffering from lower resolution, this approach would offer several practical advantages over electron microscopy. For example, treatment of BMC samples for electron microscopy can lead to deformation or alteration of structural morphology.^[26]^ In contrast, BMCs and BMC proteins imaged by confocal microscopy can remain in aqueous solution under standard buffer conditions. Also, multiple fluorophores can be utilized in confocal microscopy, thereby allowing for simultaneous imaging of multiple BMC factors. This would allow for investigation of the interactions of BMC proteins.

The BMC-H protein, PduA, which is from the *Salmonella enterica* propanediol utilization (Pdu) BMC, has previously been reported to form nanotubes when purified to homogeneity. ^19, 21, 23, 24^ To test whether PduA nanotubes could be imaged using confocal microscopy, we purified a conserved S93C mutant of PduA (referred to henceforth as just PduA). The S93C mutation has been previously shown to enable site-specific modification of PduA with a maleimide-conjugated fluorophore.^27^ PduA was successfully purified to homogeneity and fluorescently labeled, as determined by SDS-PAGE (Figure S1). TEM analysis of purified, labeled PduA confirmed its previously reported ability to form a series of superstructures including nanotubes and sheets (Figure 1a). Interestingly, some nanotubes are found in clusters about a central site (Figure 1a, arrows). In contrast, we did not observe bundles of PduA nanotubes, as was previously reported. This difference may stem from the use of different mutants of PduA and buffer conditions, a critical parameter, as compared to previous reports.^21^

**Figure 1:**
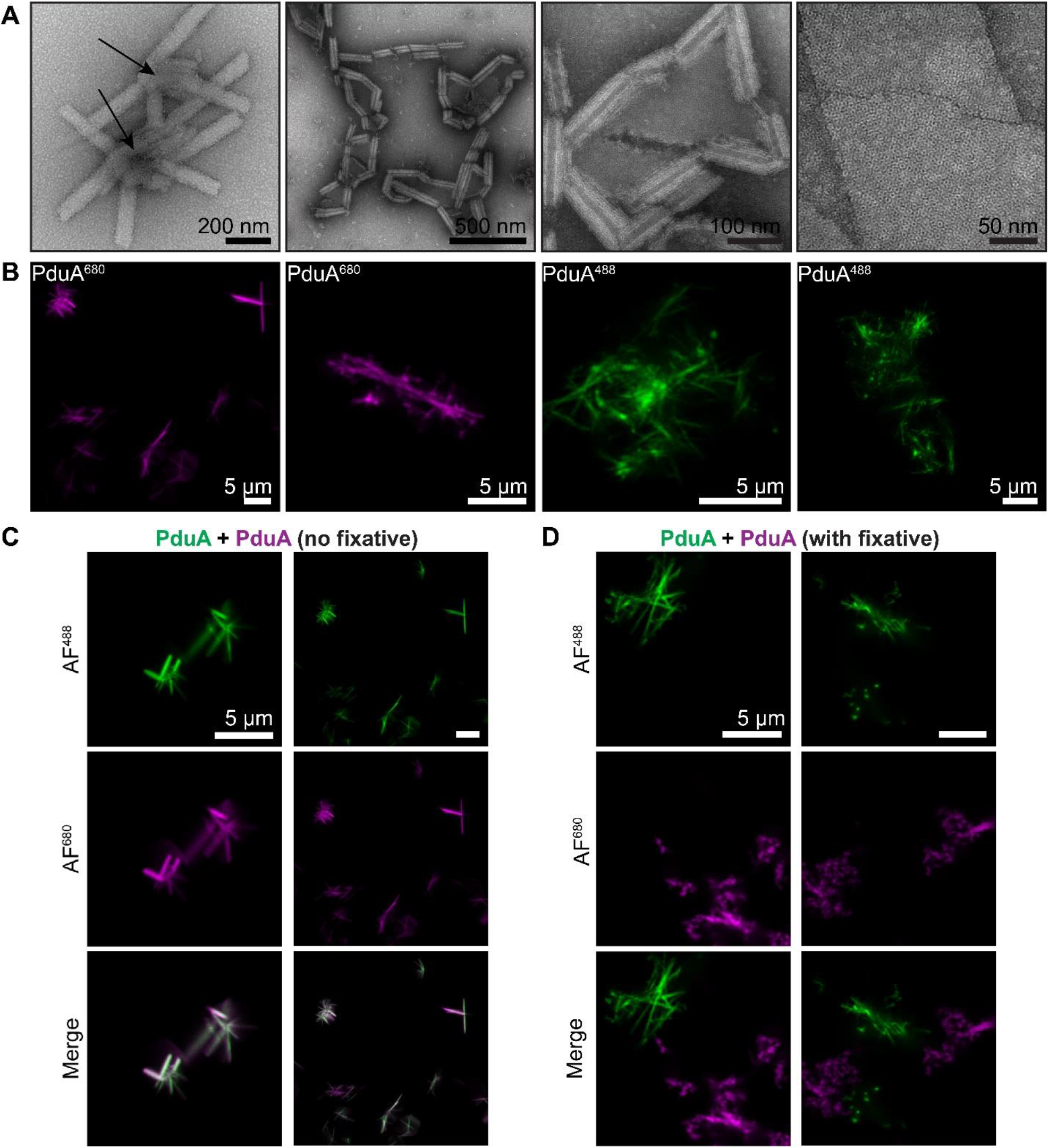
Imaging of PduA superstructures. (A) Transmission electron microscopy (TEM) analysis of PduA structures reveals clusters of nanotubes.Arrows denote clustering centers. (B) Laser scanning confocal microscopy analysis of Alexa Fluor 680-tagged PduA reveals protein nanotubes. (C) PduA samples labeled with different fluorophores were mixed together and equilibration to observe their colocalization. (D) A similar set of aliquots was fixed with paraformaldehyde prior to incubation, preventing their colocalization. All scale bars in confocal images are 5 µM.

Next, we turned to imaging the PduA structures with a confocal microscope equipped with an Airyscan detector for enhanced resolution.^28^ Under these conditions, PduA presents in dispersed clusters of objects that appear, roughly, like nanotubes (Figure 1b). These clusters are several micrometers in length and could also be observed under brightfield conditions (Figure S2). We find that these structures are agnostic to the fluorescent modification; filamentous structures were observed in samples labeled with both Alexa Fluor 680 and 488 (AF^680^ and AF^488^, respectively). More detail could be resolved with AF^488^ labeled samples due to the shorter wavelengths that are used for imaging. These results show that BMC-H nanotubes can be studied *in vitro* with a confocal microscope.

Confocal analysis can discriminate between different objects if they are labeled with different fluorophores. Given this, we sought to explore if separate sets of PduA would colocalize if equilibrated.

To investigate this, separate aliquots of AF^488^- and AF^680^-labeled PduA were mixed and equilibrated for 30 minutes. A control where the aliquots were fixed with paraformaldehyde prior to mixing was also analyzed. Confocal images were then taken at the end of the equilibration period. This revealed evidence of colocalization for the two fluorophores, suggesting that PduA superstructures may equilibrate subunits (Figure 1c). In contrast, fluorescent signals for fixed samples remained separate (Figure 1d). These results suggest that individual PduA hexamers may equilibrate between soluble hexamer and bound nanotube forms. Fixation prevents this equilibration from occurring, resulting in no exchange between separate superstructures. While we only demonstrate end-point results, these data suggest that laser scanning confocal microscopy can indeed be utilized as an experimental approach for real-time imaging of nanotubes.

### The hexamer PduJ forms distinct nanotubes

The PduJ shell protein has also been recently reported to form nanotubes.^23^ The sequence of PduJ closely resembles that of PduA, with a high degree of sequence identity across the length of the proteins. ^29^ PduA and PduJ behave similarly enough that many consider them to be interchangeable, redundant, and functionally identical to one another.^23, 29, 30^ Therefore, we sought to determine whether PduJ could form nanotubes akin to PduA. However, while our data revealed that fluorophore-labeled PduJ indeed formed nanotubes, these structures exhibited morphologies distinct from PduA. Under our assay conditions, PduJ presented as large, rosette-like structures, rather than individual nanotubes (Figure 2a, S2). Based on this observation, we speculate that the PduJ rosettes might have a complex internal structure. Like PduA, PduJ can also form sheet structures (Figure 2a, right panel). Interestingly, the higher resolution offered by TEM presented us with images that appeared to suggest that the rosettes consist of a network of flexible, interweaving nanotubes (Figure 2b). This flexibility has not previously been noted for BMC-H nanotubes. Together, these data show that PduJ can form nanotubes, but that these structures are different from PduA in terms of flexibility despite the close sequence homology of these proteins. From these data alone, we cannot offer a mechanistic explanation for the structural differences presented here, although we note that confocal microscopy offers a straightforward approach for investigating superstructures formed by PduA/J proteins containing site-directed mutations.

**Figure 2:**
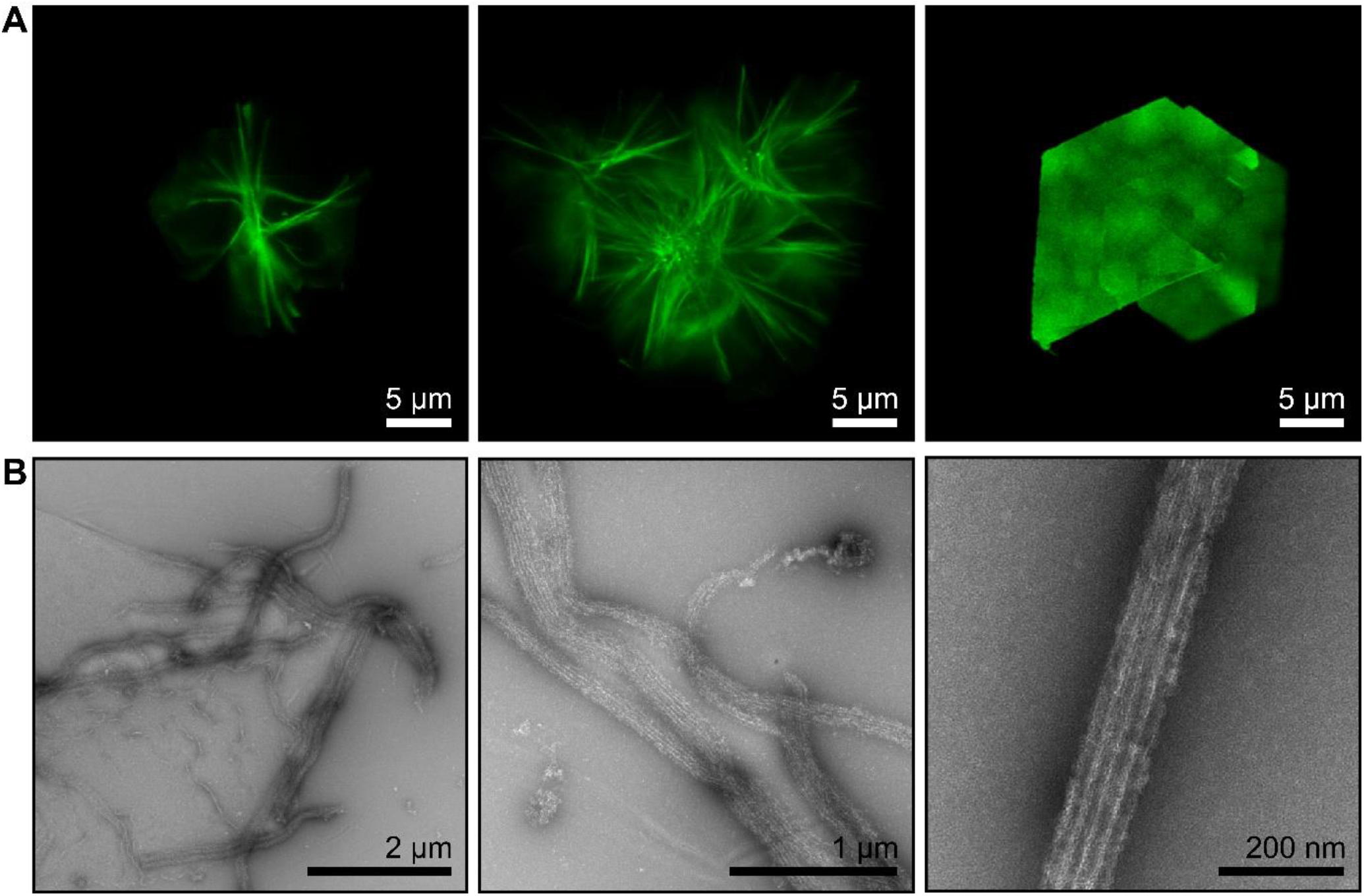
Imaging of PduJ superstructures. (A) Laser scanning confocal microscopy analysis of Alexa Fluor 488-tagged PduJ reveals protein rosettes of nanotubes and large sheet structures. (B) Transmission electron microscopy (TEM) of PduJ shows rosettes are likley to be interweaving clusters of flexible nanotubes.

### Different shell proteins can colocalize

We reasoned that confocal microscopy could provide valuable insight into the solution behaviors of the PduA and PduJ structures if different shell proteins, carrying different fluorescent tags, were equilibrated and observed as in Figure 1c and 1d. This would, for example, allow us to investigate whether different nanotubes might be able to measurably exchange protein subunits. For these experiments, we also purified the major BMC-T proteins PduB and PduB’ (an alternative translation product of the *pduB* gene that lacks the N-terminal 37 residues). After their purification, we found that PduB and PduB’ both produce aggregates in solution (Figure S3). However, PduB forms significantly larger aggregates than PduB’, suggesting that the N-terminal sequence enhances the propensity to aggregate. Previously reported PduB nanotubes were not observed under our assay conditons.^21^

PduA and PduJ, each carrying a distinct fluorescent tag, were mixed, and equilibrated on ice for 30 minutes prior to imaging. Confocal analysis revealed that subunits of PduA and PduJ structures could colocalize, albeit with interesting caveats. For instance, imaging of rosette structures, typically associated with just PduJ, showed an interesting molecular phenotype for the mixtures of PduA and PduJ. We found that the PduA/PduJ structures were hybrid, displaying a PduJ core with a periphery that featured PduA subunits (Figure 3a). From this we speculate that PduA might equilibrate initially with the exterior of the PduJ rosettes before permeabilizing inward into the PduJ superstructure. Importantly, these results reflect that not only do PduA and PduJ interact, but that some or all their respective structures are likely to be in equilibrium.

**Figure 3:**
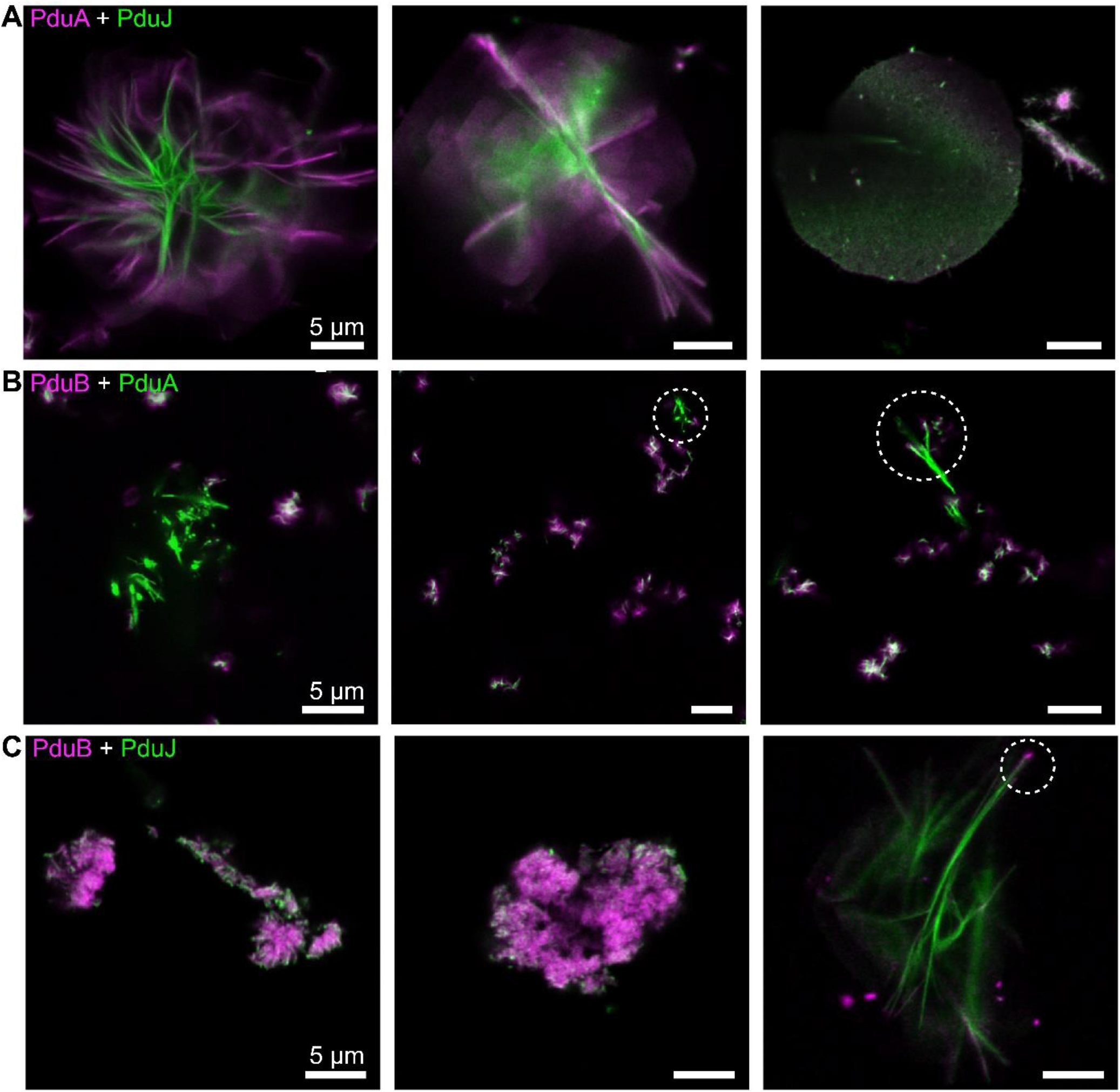
Confocal microscopy imaging of mixtures of shell proteins reveals exchange of subunits. (A) Alexa Fluor 680-tagged PduA was equilibrated with Alexa Fluor 488-labeled PduJ and then imaged by laser scanning confocal microscopy. Both rosettes and sheets were identified by microscopy; they featured a colocalized mixture of PduA and PduJ proteins. (B) Confocal microscopy imaging of PduA mixed with PduB. PduA nanotubes exhibited colocalization of PduB at or near their poles. PduA showed general colocalization with PduB aggregates. (C) Confocal microscopy imaging of PduJ mixed with PduB. PduJ appears to associate with the periphery of PduB aggregates, while PduB appears to associate at the poles of PduJ nanotubes. Circles indicate colocalization of PduB to nanotube poles. All scale bars are 5 µm.

Next, we tested whether we could observe interactions between PduA and PduJ with PduB and PduB’. PduB, and by extension PduB’, have been previously suggested to interact with PduA ^19, 31^ and, more recently, PduJ.^27^ Therefore, we again employed confocal imaging to directly investigate colocalization of PduB/PduB’ proteins with PduA and PduJ superstructures. This showed that mixing PduA and PduB led to a combination of their two independent morphologies (i.e., nanotubes plus aggregates). However, the colocalization trends differed between the PduA nanotubes and PduB aggregates (Figure 3b). PduA nanotubes displayed nearly no overlap with PduB signal except for at the end of nanotubes (circles). Conversely, it was not uncommon for PduB aggregates to display colocalization with PduA signal, although further work is needed before quantitative statements can be made about putative PduB-PduA interactions. We observed similar findings for equilibration of PduB with PduJ (Figure 3c) as well as for PduB’ (Figure S3c, d). Together, these results tentatively confirm that PduB interacts with both PduA and PduJ and suggest that individual protein subunits might be exchanged in characteristic ways for the respective PduABB’J structures. Although more work is required for quantitative statements, our data also suggests that PduB can only colocalize to the ends of PduAJ nanotubes rather than extend from them, as we observed with the PduAJ mixtures.

### Pdu BMCs can colocalize in vitro

In prior published experiments, we purified fully-formed Pdu BMCs and showed that they can be reacted with different chemical probes (*i*.*e*., fluorophores and crosslinkers).^27^ Therefore, we again purified Pdu BMCs and tagged aliquots with different fluorophores (Figure S4a). TEM analysis of treated samples confirmed they maintained similar morphology to untreated Pdu BMCs (Figure S4b), suggesting minimal sample damage had been incurred by the chemical probes. We then mixed equimolar amounts of the two BMC solutions and allowed them to equilibrate (Figure 4a). After aliquots of these mixtures were removed at short and longer time points, they were fixed with paraformaldehyde and crosslinker, and then analyzed by laser scanning confocal microscopy.

**Figure 4:**
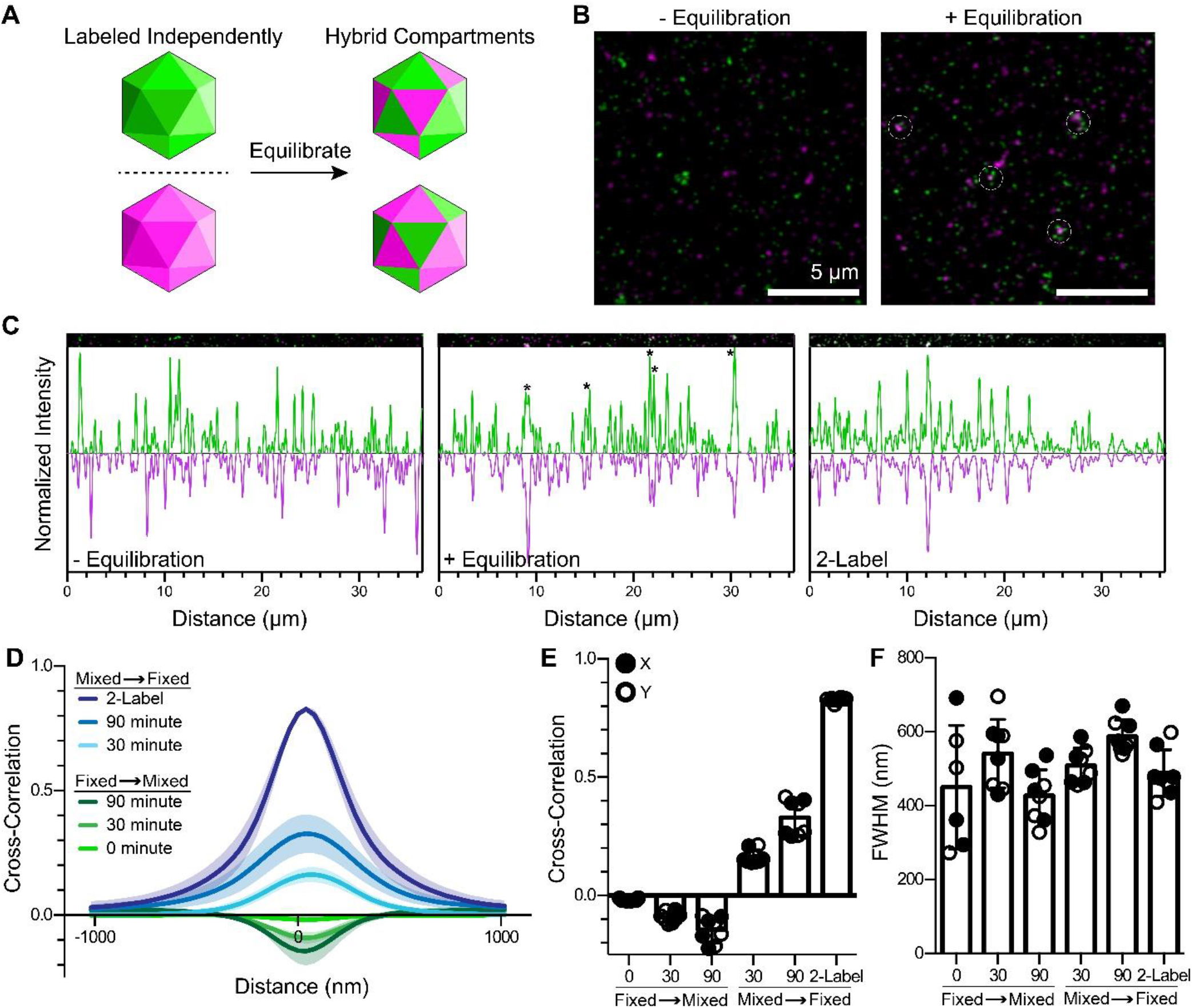
Proteins from Pdu BMCs can colocalize after equilibration. (A) Schematic of experimental design. Separately labeled BMC batches (Alexa Fluor 680, Alex Fluor 488) were mixed, equilibrated and imaged by laser scanning confocal microscopy. (B) To examine whether BMC preparations exhibited colocalization of BMC factors, samples were either imaged by confocal microscopy without being equilibrated or were equilibrated for 30 minutes on ice beforebeing imaged. Equilibrated samplesrevealed the presence of white foci (circled). Scale bars, 5 µM. (C) Plotting signal intensity from slices of micrographs. Colocalization events could be detected for equilibrated samples and for positive control samples that had been labeled with both fluorophores. (D) Pdu BMCs were either fixed prior to (“Fixed → Mixed”) or after (“Mixed → Fixed”) being allowed to equilibrate, then imaged and analyzed for cross-correlation via the van Steensel method.^[32]^ Four micrographs for each sample were analyzed in both the X and Y directions. (E) The peaks of each curve in (D) are shown in this graph. Data analyzed in the individual X and Y directions are shown. (F) The full-width of the Gaussian distributions in (C) at half-max are plotted here, showing that particle size estimates appear approximately equal for all samples.

We began by comparing differently labeled batches of Pdu BMCs that were either fixed prior to equilibration and imaging (as a negative control) or were allowed to equilibrate prior to imaging (Figure 4b). The samples that were fixed prior to being mixed appeared as distinct, distinguishable foci (shown as either green or magenta), showing no visual evidence of colocalization, likely owing to the high extent of fixation achieved (Figure S4c). In contrast, the experimental samples that were mixed and equilibrated prior to addition of the fixative showed increasing evidence of colocalization over time. As a positive control for colocalization, we also prepared a sample of BMCs that was deliberately dual-labeled by both fluorophores. To analyze these data, we quantified slices of micrographs from non-equilibrated, equilibrated, and dual-labeled samples and independently plotted the intensity of the green and magenta signals (Figure 4c). The samples that were fixed before being mixed showed an apparent random peak distribution, showing no evidence of colocalization. In contrast, the dual-labeled sample, as anticipated, showed a high degree of overlap between signal peaks, signifying colocalization of the two fluorophores. Importantly, the samples that were mixed and equilibrated showed colocalization of the two types of signal peaks.

We next turned to van Steensel cross-correlation analysis to gather deeper insight into foci colocalization in a less biased manner. Here, cross-correlation is measured as a function of a pixel shift (*i*.*e*., distance) between the green and magenta scans.^32, 33^ The resulting Gaussian curve can be positive (indicating correlation), near-zero (no correlation), or negative (anti-correlation). Pdu BMC samples that were fixed then equilibrated for either 0, 30, or 90 minutes prior to imaging all displayed modest anti-correlation (Figure 4d, 4e). The dual-labeled BMC positive control, in contrast, displayed extremely high cross-correlation of 0.85. Samples that were equilibrated for 30 or 90 minutes prior to imaging showed positive cross-correlations of 0.18 and 0.33, respectively. Van Steensel analysis also yields data on particle size (the full-width at half-maximum for the curve), which was found to be similar in all samples, thus indicating that sample treatment is not majorly disruptive (Figure 4f), in agreement with TEM results (Figure S4b). Together, these data support the hypothesis that fully formed, native Pdu BMCs can be examined with laser scanning confocal microscopy and, furthermore, may be capable of exchanging some of their components.

### Pdu BMCs interact with shell protein superstructures

The results reported herein have so far suggested that both the superstructures of Pdu shell proteins and the fully formed Pdu BMCs are capable of exchanging protein constituents. We speculate that this exchange occurs through an equilibrium that is established between the complete nanotubes/polyhedra and their individual shell proteins in solution. Here, Pdu BMCs can be roughly modeled as a specific collection of shell elements much like the nanotubes. Accordingly, we reasoned that mixing the protein nanotubes with Pdu BMCs might also result in exchange of protein subunits or even hybrid structures, which could again be observable via confocal microscopy. We first tested this by equilibrating labeled PduA nanotubes with labeled Pdu BMCs and examining the mixtures using laser scanning confocal microscopy (Figure 5a). Unexpectedly, this revealed striking images of PduA nanotubes featuring BMC foci dotted along their surface (Figure 5a). Similarly, PduA sheets also appeared to be speckled by BMC foci. To investigate the BMC foci further, we analyzed the PduA-BMC mixtures by TEM. This revealed the presence of large BMC-like entities annealed to PduA nanotubes (Figure 5b). However, these BMC structures appeared to be much less morphologically defined than in the absence of nanotubes (Figure 5b, right panel). In fact, the BMC structures appeared to envelop the PduA nanotubes in a manner that leaves the BMC entities partially deformed (Figure 5b). Similarly, we found that Pdu BMC also adhered to the exterior of the PduJ rosettes in solution (Figure 5c), although this was not investigated further with TEM.

**Figure 5:**
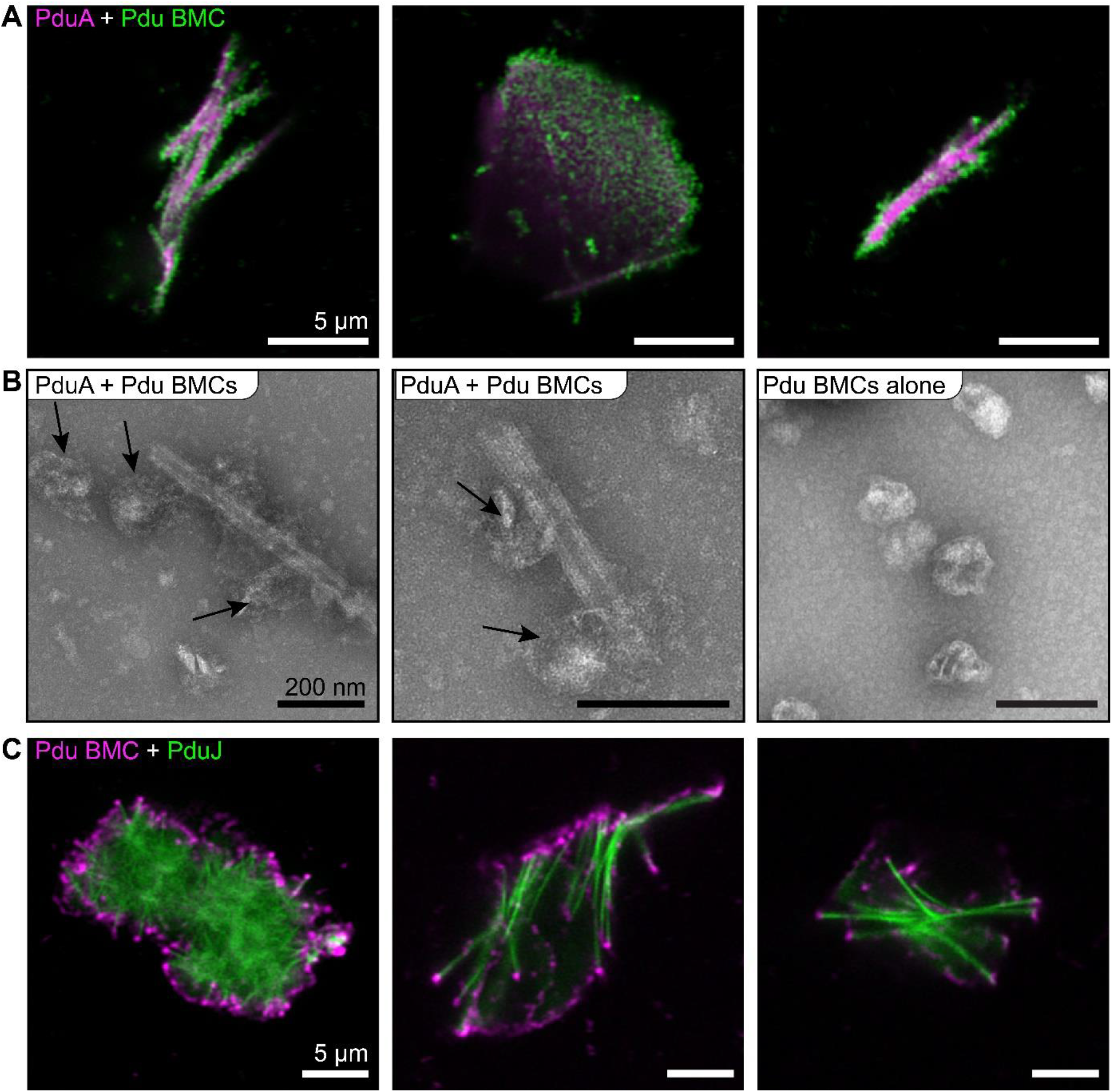
Full-sized, native Pdu BMCs interact with shell protein superstructures. (A) Alexa Fluor 680-tagged PduA nanotubes were equilibrated for 30 minutes on ice with Alexa Fluor 488-tagged Pdu BMCs, then imaged by laser scanning confocal microscopy. BMCs appear to colocalize along PduA superstructures. Scale bars, 5 µm. (B) Samples from (A) were analyzed by transmission electron microscopy (TEM). BMCs (arrows) stick to and encompass PduA nanotubes but appear to show loss of overall structural integrity. Pdu BMCs (right panel) serve as a morphology control. Scale bars, 200 nm. Alexa Fluor 488-tagged PduJ was equilibrated for 30 minutes on ice with Alexa Fluor 680-tagged Pdu BMCs, then imaged by laser scanning confocal microscopy. BMCs appear to colocalize with PduJ superstructures in characteristic ways. Scale bars, 5 µm.

### Encapsulation peptides display a preference for shell proteins

To further demonstrate the potential utility of confocal microscopy for analysis of BMC proteins, we opted to directly visualize the interactions of shell superstructures with encapsulation peptides (EPs). Cargo proteins are generally thought to interact with shell proteins and each other via short-helical termini called EPs. In the Pdu BMC system, the EP for the cargo proteins PduD and PduP have both been proposed to interact with PduA, possibly at its C-terminus.^34, 35^ Furthermore, the EP of PduP has also been suggested to interact with another BMC-H, PduK.^36^ In separate studies, the N-terminus of PduB has been speculated to act as an anchor between the inner cargo and shell. ^27, 37^ If true, we reasoned that purified EPs may associate directly with some or all shell protein superstructures, which could be observed as colocalization events.

The EPs for PduD and PduP (PduD^EP^ and PduP^EP^, respectively) were synthetically produced with an N-terminal fluorescein and C-terminal amidation to mimic their natural charge state. Both were confirmed by circular dichroism to exhibit characteristic α-helical character in solution (Figure S5). In pairwise experiments, we equilibrated the peptides individually with AF^680^ labeled shell proteins (PduABB’J) prior to imaging by confocal microscopy. This revealed strong colocalization with PduD^EP^ and PduP^EP^ to PduB but not PduB’ (Figure 6a, b). From this, we speculate that the N-terminal portion of PduB may facilitate inwards-facing cargo interactions. This is consistent with recent crosslinking mass spectrometry experimentation, which provided direct evidence for interactions between PduB and numerous EP-containing cargo proteins in a native context, as well as other data suggesting PduB is a shell-cargo anchor point.^27, 37^ PduA nanotubes were similarly found to colocalize with PduD^EP^ and PduP^EP^ (Figure 6a). In contrast, colocalization with PduJ was observable but comparatively reduced (Figure 6d). EPs did not appear to qualitatively influence shell protein morphologies and did not form structures on their own under the conditions used in our experimentation. Together, these data tentatively support an interaction between encapsulation peptides and the N-terminus of PduB and between the EPs and PduA. These data are consistent with the hypothesis that PduB and PduA are likely to be primary factors in facilitating shell-cargo interactions.

**Figure 6:**
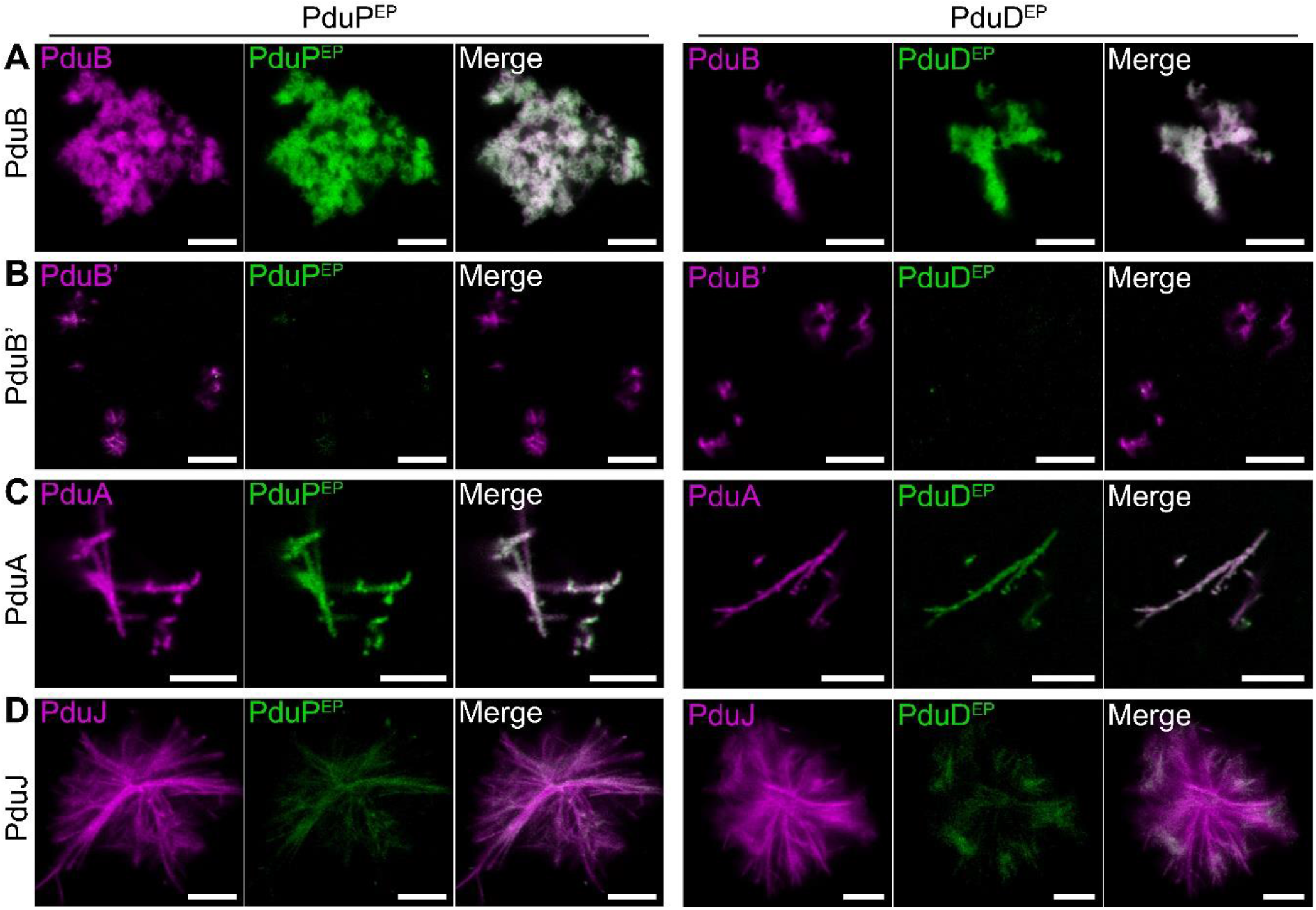
Association of encapsulation peptides with shell protein superstructures. Purified, synthetic, fluorocein-tagged encapsulation peptides (EP) for PduP (PduP^EP^) and PduD (PduD^EP^) were equilibrated with either (A) PduB, (B) PduB’, (C) PduA, or (D) PduJ. In all instances, samples were equilibrated for 30 minutes on ice before being imaged by laser scanning confocal microscopy. Both EPs behaved similarly; a greater degree of colocalization was observed between the EPs and PduB and PduA. All scale bars are 5 µm.

## Discussion

In total, the data presented herein suggests that confocal microscopy can be used to investigate exchange of subunits between shell protein superstructures and between fully-formed BMCs. Here, we imagine the Pdu BMC as being in an equilibrium between complete Pdu BMCs and individual shell elements in solution (Figure 7a). Interestingly, previously published atomic force microscopy experiments showed that BMC-H sheets assemble rapidly and can exchange subunits;^38,39^ perhaps BMC polyhedra facets behave similarly. We speculate that BMCs may exist along a continuum that is an evolved adaptation to suit different needs. Carboxysomes, for instance, may exist on the more rigid side of this scale, maintaining a hard shell (Figure 7b). They have been previously noted to formmuch moreregular polyhedral structures than metabolosomes. Furthermore, carboxysomes can survive direct inheritance through generations^40^ and directly inhibit the passage of small molecules like O_2_,^41, 42^ which would otherwise inhibit carbon fixation^43, 44^ - features that are consistent with a rigid, relatively static shell structure. In contrast, the Pdu BMC has a noticeably less regular structure and will permit the passage of relatively large chemicals.^27^ While the inner metabolism of some BMCs may create toxic aldehydes that the shell sequesters,^45, 46^ this does not appear to be a necessity as these metabolic pathways are sometimes encoded without need for shell proteins.^47^ Furthermore, bacteria which encode multiple different BMCs heavily regulate them as to prevent their intermixing.^48^ It may be the case that metabolosomes are less rigid in order to be more responsive to environmental changes, perhaps so that they can be rapidly assembled and disassembled based on the cell’s metabolic needs, while, in contrast, carboxysomes fulfill a more long term, and fixed, metabolic requirement. Perhaps then, the shell layer of BMCs may exhibit a high enough degree of protein exchange that it can be measured experimentally, and may therefore be reflected in the colocalization of BMC proteins observed in this study. Although it was not investigated in this study, we suspect that carboxysomes will exhibit less colocalization of their shell proteins, given the expectation that they might form more rigid shell structures. However, the data presented herein is a largely qualitative assessment of this phenomena. More evidence, particularly from *in vivo* systems, is needed to clarify the role of BMC protein intermixing. Indeed, an enhanced understanding of BMC dynamism would strengthen how we view the BMC life cycle from biogenesis to degradation.

**Figure 7:**
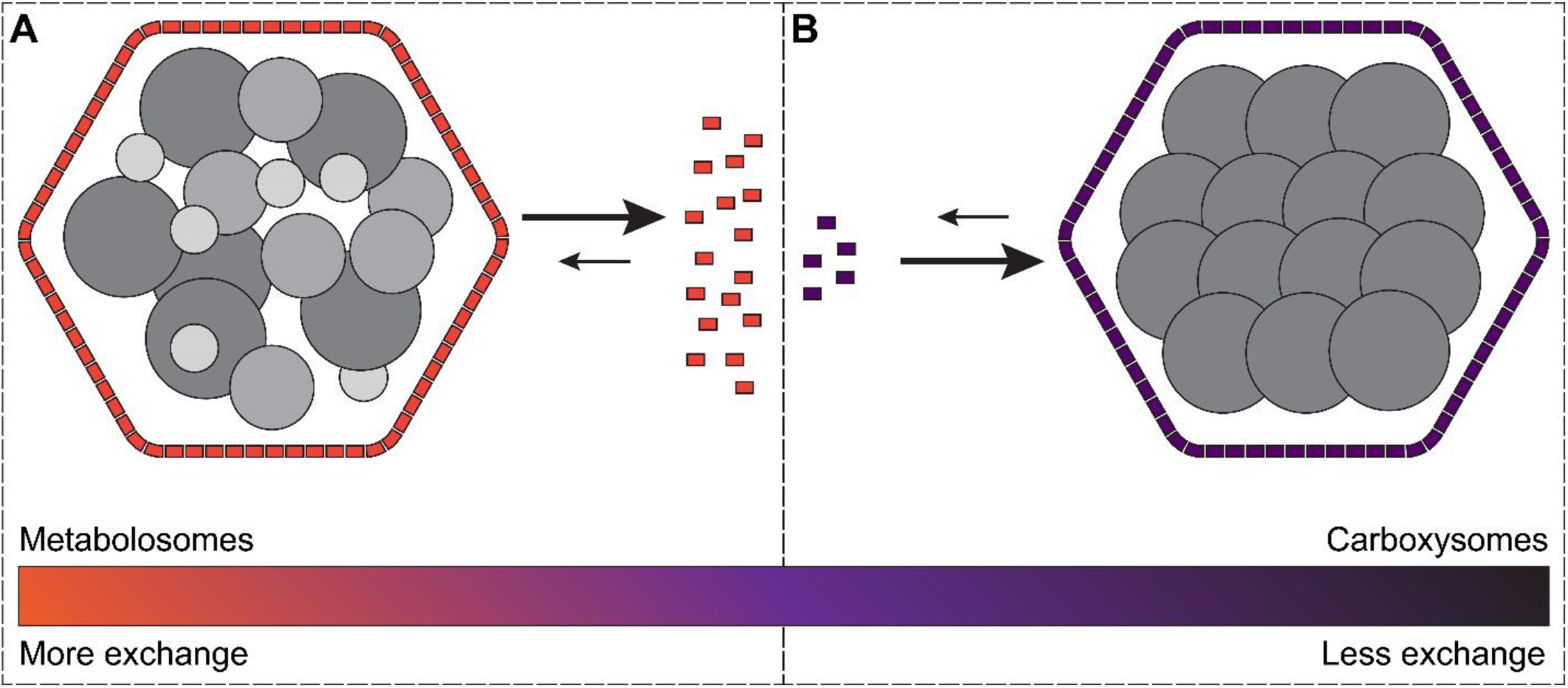
Models for the behavior of the Pdu BMC. (A) Metabolosomes, like the Pdu BMC studied here, may be capable of more subunit exchange relative to (B) carboxysomes, which are known to exhibit rigid structures. If the shell layers are exchangable, the BMCs may exist in equilibrium with cytoplasmic shell proteins.

The fact that the PduA and PduJ structures display differences calls into question the assumption that PduA and PduJ are functional redundant for shell assembly. A recent report suggested that the PduB trimer specifically accommodated PduA and PduJ subunits along different interaction interfaces. ^27^ Together, these data suggest that assembly of the shell layer might be affected by subtle differences in BMC-H proteins, which are particularly widespread among BMC operons. ^3^ Elucidating these subtle differences among BMC-H will help future efforts at designing novel BMC shells of various sizes and packaging capacity.

Colocalization analysis also revealed that several Pdu EPs interacted specifically with the shell proteins PduAB but did not appear to associate with PduB’J under our assay conditions. Curiously, PduAB are also encoded by the first two genes in the *S. enterica pdu* operon.^49^ This observation may be consistent with the speculation that PduAB may colocalize with cargo to an EP-driven pro-metabolosome in the initial stages of biogenesis coupling together shell and cargo. PduAB could then seed further shell envelopment and eventually lead to mature BMCs. This model is analogous to the concomitant assembly proposed for α-carboxysomes,^1, 50^ despite metabolosomes being more related to β-carboxysomes.^51^ We speculate that a liquid-liquid phase separation (LLPS) mechanism is likely to apply to the early stages of metabolosome assembly, as supported by recent reports focusing on the Pdu BMC.^52^ This has also been suggested for both α- and β-carboxysomes too, supporting the notion that LLPS may be a common mechanism for BMC biogenesis.^53, 54^ Our data does not support the shell or shell protein structures themselves being liquid. In conjunction with other reports that mainly focus on the condensation of cargo, perhaps the shell participates in early BMC biogenesis but then acts to constrain the inner coacervate in a mature BMC.

Confocal microscopy is a powerful tool to study both *in vitro* and *in vivo* colocalization events. Here, we leveraged its ability to differentiate protein components to begin to understand how BMC shell protein structures and whole BMC polyhedra interact in solution. This experimental approach complements traditional TEM approaches. We anticipate that the experimental conditions for laser scanning confocal microscopy can be varied extensively to ask many different questions on BMC or the superstructures formed by BMC shell proteins. For example, future studies may wish to go further to include time-lapsed measurements, or to incorporate emerging super-resolution microscopy techniques or fluorescence correlation spectroscopy.^55^ Therefore, we expect confocal microscopy will continue to serve as an important tool for investigating shell protein nanotubes in solution, as we showcase here, as well as how they might operate as synthetic *in vivo* scaffolds or if developed as potential biomaterials.

## Acknowledgements

We thank Dr. Amy Beaven of the University of Maryland Imaging Core for training and expertise in laser scanning confocal microscopy. Purchase of the Zeiss LSM 980 Airyscan 2 was supported by award Number 1S10D025223-01A1 from the National Institute of Health. We also thank Dr. Tim Miguel of the University of Maryland Laboratory of Biological Ultrastructure for his help with transmission electron microscopy imaging. This work was initiated using funding to W.C.W. from NIH/NIAID R01AI110432.

## Contributions

D.S.T and W.C.W designed the study; D.S.T performed research; D.S.T. analyzed the data; and D.S.T and W.C.W wrote the paper.

## Methods

### Purification of shell proteins

All shell proteins were purified using the same set of conditions. Vectors encoding for the shell pro teins PduABB’JN were transformed into T7 Express E. coli. For protein production, strains were streaked from glycerol stocks onto LB agar with carbenicillin. Single colonies were then used to produce overnight starter cultures. The next day, starter cultures were used to inoculate 500 mL of 2xYT broth supplemented with carbenicillin and grown to at OD600 of 0.6-0.8. Once at the appropriate density, IPTG was added to 0.5 mM to induce protein expression. Cultures were further incubated at 37°C for 3 hours bef ore being pelleted and frozen at −80°C. Pellets were then thawed, resuspended in 40 mL Buffer A (50 mM Tris-HCl pH 8.0, 1 M NaCl, 5 mM DTT, 10 mM imidazole) supplemented with 5 mM MgCl_2_, 1 U/mL Baseline-Zero DNase (Lucigen), and 0.5 mM PMSF. Cells were then lysed with sonication and insoluble matter was pelleted by centrifugation at 13,000 x g for 20 minutes. Soluble matter was then incubated with 1 mL of nickel-NTA affinity beads that had been pre-equilibrated in Buffer A for 30 minutes on ice. The flowthrough was drained and beads were washed with 5 mL of Buffer A followed by 5 mL of Buffer B (50 mM Tris-HCl pH 7.4, 1 M NaCl, 35 mM imidazole). This second wash removes non-specifically bound proteins and equilibrates bound samples for fluorescent labeling. The appropriate fluorophore (either Alexa Fluor maleimide 488 or 680) was dissolved in 1 mL of Buffer B and added to the bead bed where the reaction was allowed to proceed for 2 hours at 4°C. The excess fluorophore was then drained off and the beads were washed again with 2x 5 mL of Buffer B supplemented with 5 mM DTT. Labeled proteins were then eluted in Buffer C (50 mM Tris-HCl pH 8.0, 1 M NaCl, 5 mM DTT, 250 mM imidazole) and dialyzed into storage buffer (20 mM Tris-HCl pH 8.0, 100 mM NaCl, 5 mM DTT). Aliquots were dispensed and flash frozen for future use.

### Pdu BMC Purification from a heterologous host

*S. enterica* Pdu BMCs were expressed and purified from an *E. coli* host.^56^ Briefly, A single colony of R995 + PduST was used to inoculate 100 mL of 2xYT supplemented with kanamycin and 0.5% 1,2-propanediol and grown overnight at 37°C for ∼16 hours. The next day, the culture was pelleted and resuspend in 20 mL of 50 mM Tris-HCl pH 8.0, 200 mM KCl, 5 mM MgCl_2_, 5 mM β-mercaptoethanol (βME), 0.5 mM EDTA, 0.5 mg/mL lysozyme, 0.5 mM PMSF and 50% B-PER (Thermo) (v/v) and gently rocked at RT for 25 minutes and then placed on ice for 5 minutes. Genomic DNA was fragmented by 2x 1s pulses from a sonicator. Insoluble matter was pelleted for 20 minutes at 13,000 x g at 4°C. BMCs were pelleted at 17,500 x g at 4°C for 40 minutes. Pelleted BMCs were then resuspended in labeling buffer (20 mM phosphate pH 7.4, 50 mM KCl, 5 mM MgCl_2_) and pelleted again for 20 minutes. The final BMC pellet was resuspend in 1 mL of labeling buffer and stored at 4°C for less than two weeks. BMCs were quantified via Bradford Assay against a BSA standard.

### Preparing Pdu BMCs for confocal imaging

The preparation of Pdu BMCs for confocal imaging is outlined in Figure S4. Briefly, purified Pdu BMCs were diluted to 1.0 mg/mL in labeling buffer (20 mM phosphate pH 7.4, 50 mM KCl, 5 mM MgCl_2_) and labeled with 125 μM fluorophore for 15 minutes on ice. The reaction was quenched with 5 mM DTT. Pdu BMCs were then pelleted at 200,00 x g for 15 minutes at 4°C and resuspended in fresh labeling buffer to remove all excess fluorophore. When appropriate, Pdu BMCs were then fixed in the presence of 2% paraformaldehyde and 5 mM BS(PEG)5 crosslinker for 30 minutes at room temperature.

### Laser scanning confocal microscopy imaging

All samples were imaged on a Zeiss LSM 980 Airyscan at room temperature using a 63x objective with immersion oil. All samples were likewise prepared in labeling buffer (20 mM phosphate pH 7.4, 50 mM KCl, 5 mM MgCl2) for all experiments. Briefly, 20 μL of sample solution (typically 20 μM protein) was pipetted directly onto a #1.5 coverslip (Fisherbrand, 12544E). Initial focus was achieved by focusing on the edge of the sample droplet under transmission mode before switching to fluorescence and focusing down to the coverslip surface. Structures are plentiful and bright enough to allow live calibration of the Airyscan detector in super resolution mode. The fastest possible image scan time was always used with a minimum pixel scan time of 1.00 μs. All captured and reported images had standard Airyscan processing applied. Captured, unedited images were all analyzed in ImageJ. The JACoP ImageJ plugin was used to facilitate van Steensel cross-correlation analysis (Bolte and Cord; 2006).

### Transmission electron microscopy

Samples were thawed and diluted in labeling buffer (20 mM sodium phosphate pH 7.4, 50 mM KCl, 5 mM MgCl2) to 20 μM and 5 µL was applied to a formvar coated copper grid, 200 mesh (FCF200-CU, Electron Microscope Sciences). Samples were air dried for 15 minutes and excess solution was wicked off. Salt was removed by 3x 5 µL additions of deionized water before staining with 3 µL of 2% uranyl acetate for 10 seconds. Samples were imaged using a JEOL 100CXII electron microscope.

### Circular dichroism of EPs

Synthetically derived EPs were purchased from Genscript. Dried peptides were dissolved in ice-cold deionized water to a concentration of 500 μM and stored on ice until analyzed. For analysis, peptides were diluted to 50 μM in deionized water in a 1 mM quartz cuvette. Measurements were taken on a Jasco J-810 circular dichroism spectrometer from 180 to 300 nm in 1 nm increments with a scan speed of 100 nm/min in triplicate with moderate smoothing.

### EP-shell protein binding assay

Synthetic EPs were resuspended in 10 mM Tris-HCl pH 8.0. After EP resuspension, EPs and shell proteins were mixed (4 µM and 40 µM, respectively) and equilibrated on ice for 30 minutes before imaging. All images between samples were taken with the same laser intensity and detector settings.

