## Supplemental Figures for "*In Vitro* Analysis of Bacterial Microcompartments and Shell Protein Superstructures by Confocal Microscopy"

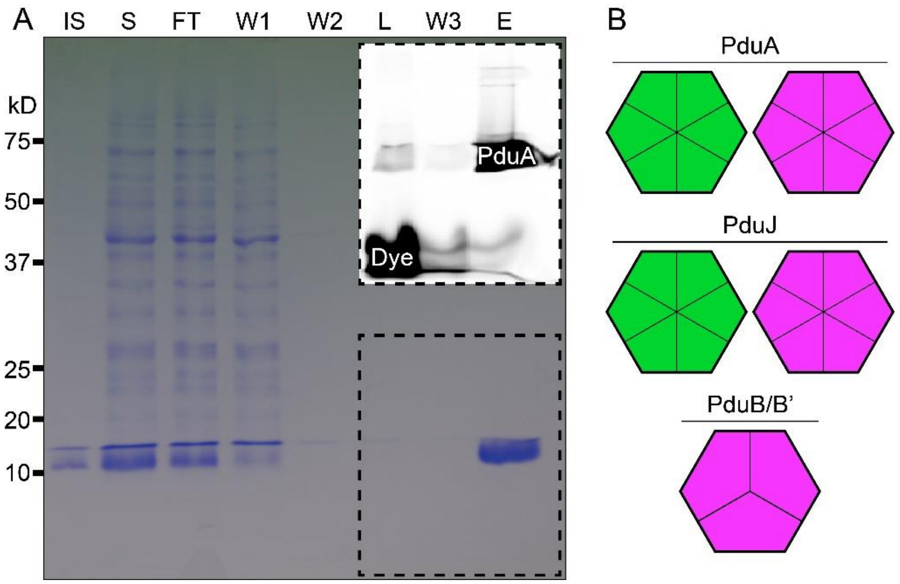


**Figure S1: Representative data showing purification and fluorophore-labeling of Pdu PduA.** (A) The purification profile for PduA S93C that was labeled with Alexa Fluor 680 maleimide. IS, insoluble fraction; S, soluble fraction; FT, flowthrough; W1, wash 1; W2, wash 2; L, eluant post-labeling; W3, wash 3; E, elution. The inset outlined in a dashed line shows the results of scanning the gel for fluorescence. This revealed the extent of $AF^{680}$ removal from the eluted protein fraction. (B) In this work, PduA, PduJ, PduB, and PduB' were purified. Proteins are represented graphically as their oligomeric states. Green represents purification with an Alexa Fluor 488 label, while purple corresponds to Alexa Fluor 680.

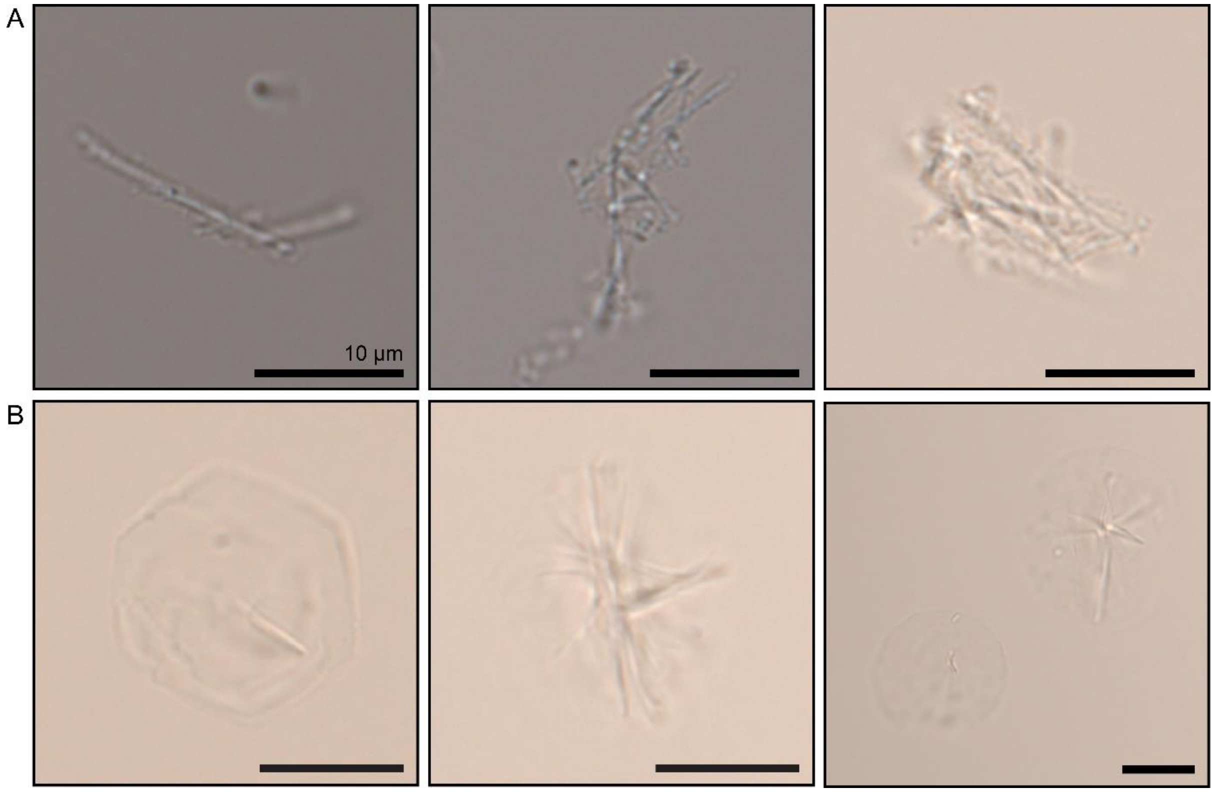


**Figure S2: Brightfield images of BMC-H assemblies.** (A) PduA was observed as nanotubes, typically in clusters and occasionally sheets. (B) PduJ was observed in rosette-like structures and highly hexagonal-like sheets. All scale bars are 10 µm.

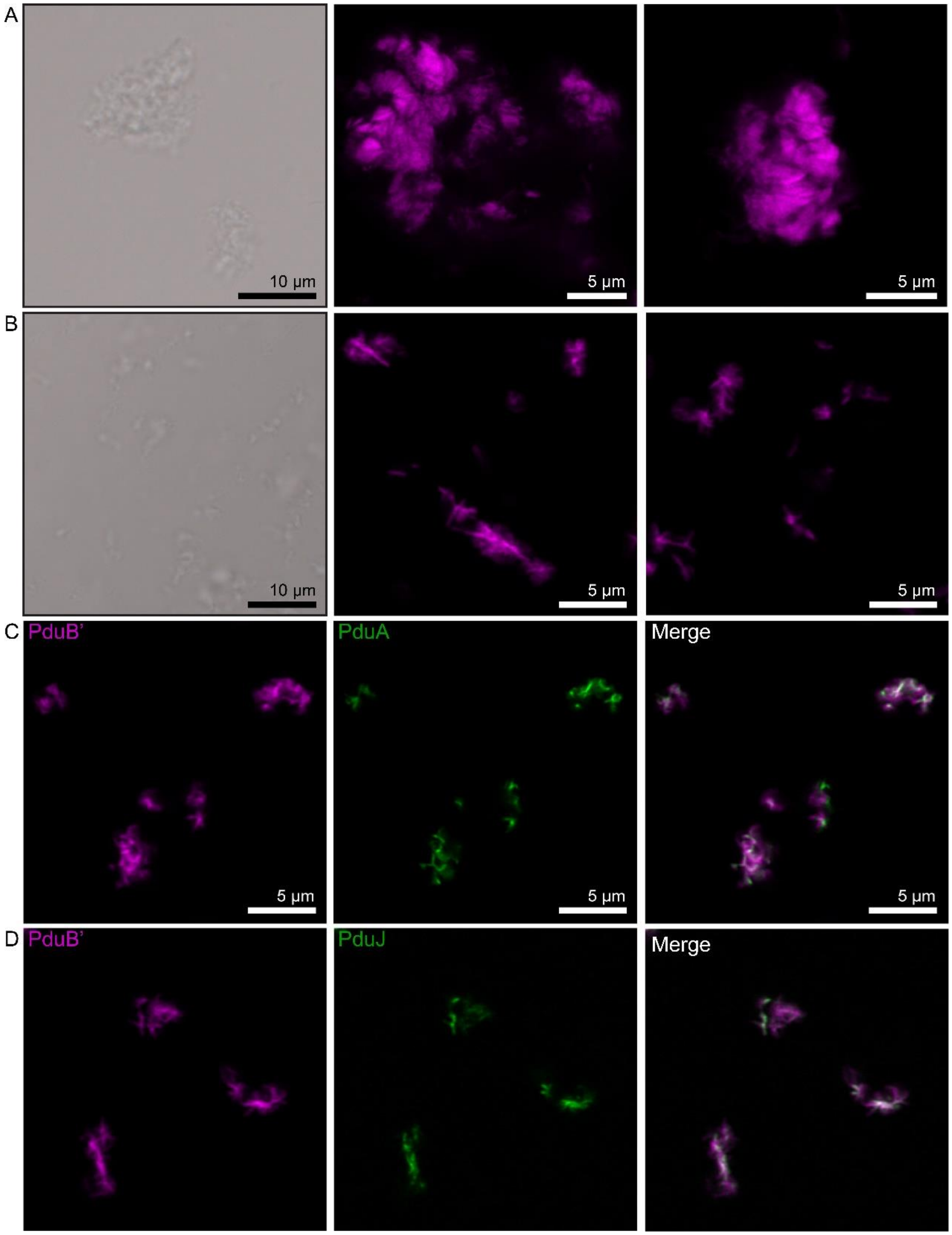


**Figure S3: The shell proteins PduBB' form aggregates.** (A) PduB and (B) PduB' differ on the basis of an additional N-terminal extension, which PduB' lacks. This extension lays on the convex, interior surface of a Pdu BMC. Both PduB (A) and PduB' (B) form protein clusters. The clusters of PduB are much larger than those of PduB'. PduB' aggregates colocalize with PduA (C) and PduJ (D) similarly to PduB.

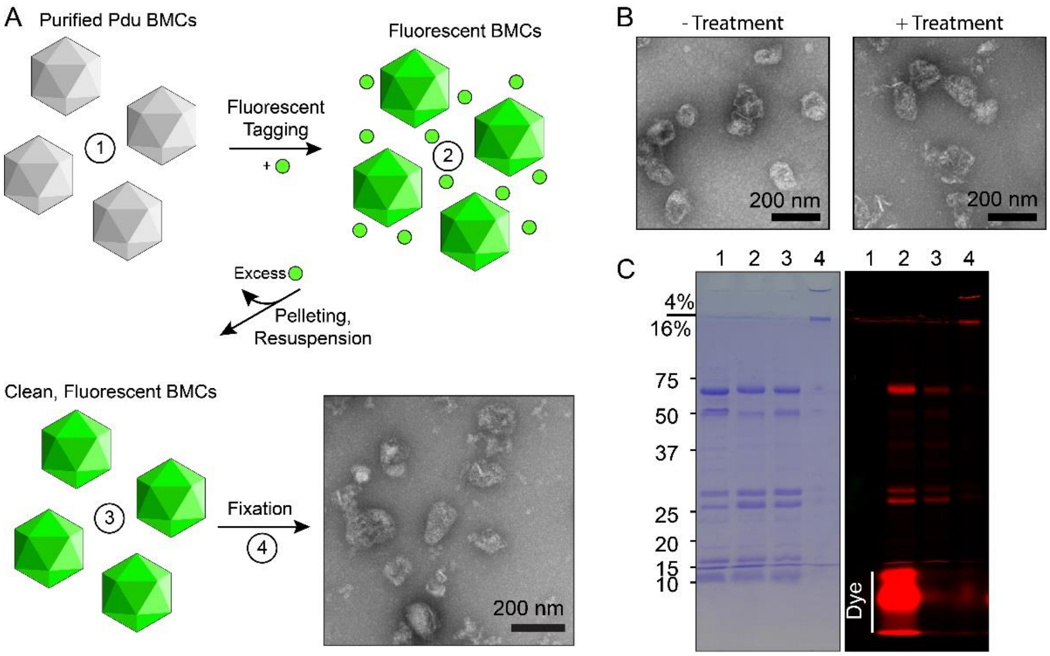


**Figure S4: The preparation of Pdu BMCs for imaging with a laser scanning confocal microscope.** (A) A scheme showing the steps in the preparation. Numbers correspond to the lanes in (C). (B) Treated samples present similarly as native Pdu BMCs when visualized by TEM. (C) Samples at various stages of preparation were resolved by SDS-PAGE to evaluate the preparation scheme. Lane 1, Purified Pdu BMCs; Lane 2, Pdu BMCs after thiol labeling with $AF^{680}$; Lane 3, Pdu BMCs after pelleting to remove excess dye left in the supernatant. Nearly all excess dye is removed; Lane 4, Pdu BMCs after fixation exhibiting band shifts. Almost no sample enters the resolving gel.

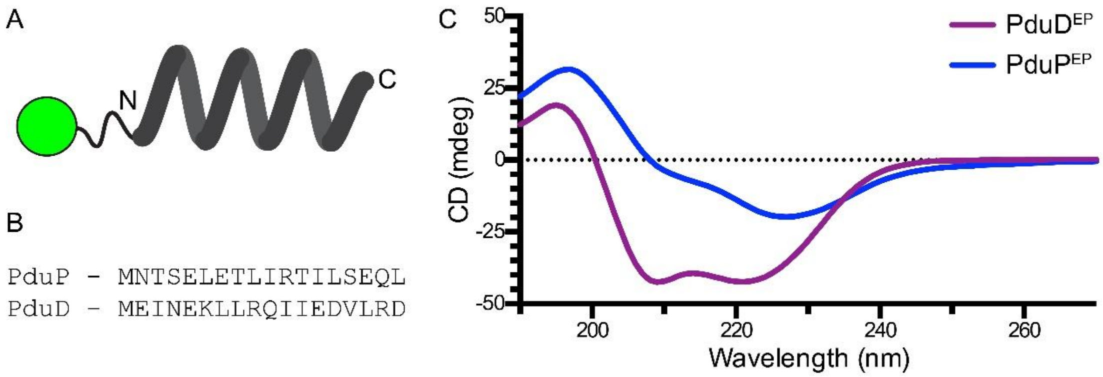


**Figure S5: The encapsulation peptides (EPs) used in this work.** (A) Cartoon schematic of the synthetic derived EPs tagged with fluorescein (green ball). (B) Sequence comparison of the EPs for PduP and PduD. (C) Circular dichroism of the selected encapsulation peptides. Both show α-helical properties, although $PduP^{EP}$ less-so. We suspect this is due to greater insolubility of $PduP^{EP}$ in pure water, where diluted $PduP^{EP}$ would form a somewhat opaque solution, compared to $PduD^{EP}$.
